# Non-canonical signaling mechanisms of short-chain fatty acid receptors in glucagon-like peptide-1 (GLP-1) releasing enteroendocrine cells

**DOI:** 10.64898/2026.03.01.708924

**Authors:** K.E. Masse, B.N. Lee, H. Wu, J. Han, P. Larraufie, F. Reimann, F.M. Gribble, V.B. Lu

**Affiliations:** Department of Physiology and Pharmacology, University of Western Ontario, London, ON, Canada; Institute of Metabolic Science and MRC Metabolic Diseases Unit, University of Cambridge Addenbrooke’s Hospital, Cambridge, UK; Children’s Health Research Institute, London Health Sciences Centre, London, Ontario, Canada

**Keywords:** FFA2, FFA3, GLP-1, SCFAs, Ketone Bodies, Enteroendocrine Cells

## Abstract

**Objectives:** Free fatty acid receptors 2 and 3 (FFA2 and FFA3) are activated by nutrient-derived metabolites such as short-chain fatty acids (SCFAs) and ketone bodies, produced by the gut microbiota and host, respectively. This study aimed to investigate the intracellular signaling pathways recruited in glucagon-like peptide-1 (GLP-1) releasing enteroendocrine cells following activation of FFA2 and FFA3 to resolve the impact of nutrient status on enteroendocrine cell function.

**Methods:** Experiments were performed using primary mouse colonic cultures and the mouse enteroendocrine cell line, GLUTag cells. Expression analysis by bulk RNA sequencing was used to determine expression of FFA2 and FFA3 in GLP-1 releasing cells. Measurement of GLP-1 secretion by sandwich ELISA was used to assess enteroendocrine cell function. Live-cell measurements of intracellular calcium and cAMP levels were performed to assess canonical second messenger signaling pathways.

**Results:** A SCFA mixture stimulated GLP-1 secretion from both primary mouse colonic cultures and GLUTag cells. In GLUTag cells, the FFA2 ligand 4-CMTB inhibited GLP-1 release independent of Ga_q_- and Ga_i_-signaling as neither YM-254890 (Ga_q_ inhibitor) nor pertussis toxin (Ga_i_- uncoupler) altered its effect. However, 4-CMTB did elevate cAMP levels, suggesting an indirect mechanism for the increase in cAMP production. Stimulation of FFA2 with the Ga_i_-biased ligand AZ1729 or the ketone body acetoacetate inhibited GLP-1 release and cAMP accumulation. AZ1729 was insensitive to pertussis toxin and OZITX, supporting atypical FFA2 signaling. Stimulation of FFA3 with AR420626 or the ketone body β-hydroxybutyrate increased GLP-1 secretion from GLUTag cells, an effect that was not mediated by cAMP production. AR420626, but not β-hydroxybutyrate increased intracellular calcium levels.

**Conclusions:** Overall, activation of FFA2 inhibited secretory function in GLP-1-releasing enteroendocrine cells, whereas activation of FFA3 stimulated GLP-1 secretion via distinct intracellular signaling mechanisms.

Graphical Abstract

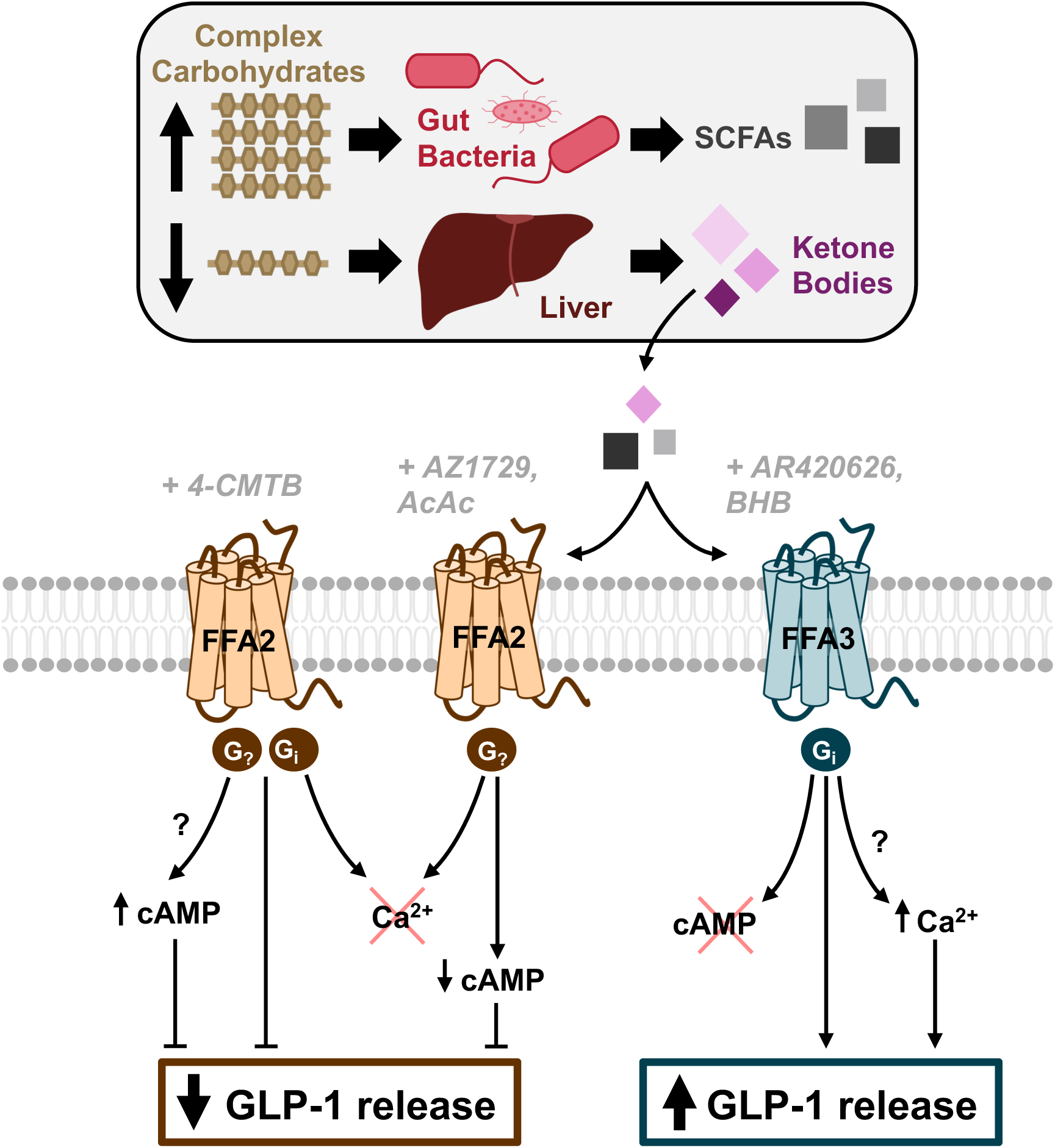

**Highlights:**

- Exposure to physiological levels of SCFAs stimulates GLP-1 secretion from colonic EECs
- FFA2 and FFA3 regulate GLP-1 release via non-canonical signaling pathways
- Ketone bodies activate SCFA receptors to differentially modulate GLP-1 levels
- Ligand bias enables nutrient-dependent tuning of EEC gut hormone secretion

## 1. Introduction

Enteroendocrine cells (EECs) are specialized gastrointestinal epithelial cells that regulate digestion and metabolism through the release of gut hormones [1]. For instance, glucagon-like peptide-1 (GLP-1) released from EECs enhances insulin secretion and slows gastric emptying to promote satiety [2]. Synthetic mimetics of GLP-1, such as semaglutide (Ozempic®/Wegovy®), have been effective in managing type 2 diabetes and obesity. Despite the concentration of GLP-1 releasing cells in the distal gut, stimulation of endogenous release of GLP-1 has not been exploited therapeutically to the same degree as exogenous GLP-1 administration, mainly due to our gap in knowledge on the precise mechanisms governing GLP-1 release in the distal gut.

Gut hormone secretion from EECs is primarily stimulated by the presence of nutrients [3], [4]. However, in the distal small intestine and colon, the direct availability of dietary nutrients to stimulate GLP-1 releasing EECs is limited. Instead, other luminal mediators, such as microbial and host-derived metabolites in circulation, likely serve as key regulatory signals for GLP-1 secretion [5], [6]. Short-chain fatty acids (SCFAs) for instance, produced by bacterial fermentation of undigested carbohydrates, are the most abundant microbial metabolites in the colon, reaching concentrations greater than 100 mM in the human caecum and ascending colon [7]. The predominant SCFAs - acetate (C2), propionate (C3), and butyrate (C4) - are generated in a molar ratio of approximately 3:1:1, respectively [7], [8]. In contrast, the chemically related ketone bodies, acetoacetate (AcAc) and β-hydroxybutyrate (BHB), are host-derived metabolites that typically circulate at lower concentrations (<0.5 mM) but can rise significantly during prolonged fasting, uncontrolled diabetes or intense physical exertion [9], [10], [11], [12].

GLP-1 releasing EECs detect nutrients through a combination of transporter-associated uptake mechanisms [13] and cell surface G-protein coupled receptors (GPCRs). SCFAs activate the free fatty acid receptors 2 and 3 (FFA2 and FFA3) [14], [15], both of which are expressed in mouse [5], [6], [16] and human [17], [18] GLP-1 releasing EECs. Although FFA2 and FFA3 share ∼40% sequence similarity and respond to SCFAs ranging from two to six carbons in length, FFA2 exhibits a higher rank order of potency for C2 and C3, whereas FFA3 preferentially binds to carbon chain lengths of C3 to C5 [14], [15], [19]. Additionally, FFA2 couples to both Ga_q_- and Ga_i_-protein families, while FFA3 exclusively activates the Ga_i_-signaling pathways [20]. BHB has been shown to inhibit neurotransmitter release via FFA3 activation in the sympathetic nervous system [21], [22] and has recently been implicated in suppressing glucose-induced GLP-1 secretion from EECs [23]. Beyond receptor activation, the signaling pathways recruited by SCFAs and ketone bodies to modulate GLP-1 release from EECs remain poorly understood. The aim of this study was to investigate the effects and signaling mechanisms downstream of activated SCFA receptors in GLP-1 releasing EECs. We found activation of FFA2 and FFA3 had opposing effects on GLP-1 release and did not recruit signaling pathways previously characterized in overexpression systems, demonstrating unique signal transduction mechanisms in endogenous GLP-1 releasing EECs. With a better understanding of these regulatory mechanisms, we can assess a reserve pool of GLP-1 stored in EECs for the prevention and treatment of metabolic disease.

## 2. Materials and methods

### 2.1. Primary murine colonic cultures

Animal studies were approved by the Western University Animal Care Committee according to guidelines established by the Canadian Council on Animal Care. Male and female C57BL/6 mice aged 2-3 months were euthanized and colonic crypts were isolated and cultured as previously described [24], [25]. Briefly, the colon was flushed thoroughly with ice-cold PBS, and the muscle layer was removed. The colon was cut open longitudinally, minced into ∼1-2 mm^2^ tissue fragments and digested with Collagenase type XI dissolved in DMEM medium (0.35 mg/mL) at 37°C and mechanical disruption. Following digestion, the suspension was filtered through a 70 µm cell strainer and plated onto 24-well plates coated with 2% Basement Membrane Extract (BME, Biotechne) for secretion experiments. The ROCK inhibitor Y-27632 (10 µM; Tocris Bioscience) was added to final cell suspensions to prevent anoikis.

### 2.2. Cell culture and transfection techniques

GLUTag cells (kind gift from Daniel Drucker, Toronto) were maintained on 0.2% Matrigel-coated flasks and low glucose (5.6 mM) DMEM supplemented with 10% FBS, 2 mM L-glutamine, 100 units/mL penicillin and 0.1 mg/mL streptomycin. Cells were passaged 2-3 times weekly and maintained at 37°C in 5% CO_2_ until passage 35. Cells for experiments were plated on 2% Matrigel-coated 24-well plates for secretions, 96-well plates for cAMP imaging, or 35 mm glass-bottom dishes (MatTek) for Ca^2+^ imaging.

For transfections, GLUTag cells were seeded onto 6-well plates at a density of 0.8-1.0 x 10^6^ cells/well. The following day, semiconfluent GLUTag cells were transiently transfected with 2 µg pcDNA3.1(+)-OZITX S1 (Addgene #184925) using Lipofectamine 3000 (Invitrogen) according to the manufacturer’s instructions. 48 h post-transfection, cells were plated for GLP-1 secretion assays. GLUTag cells stably expressing the cAMP biosensor GloSensor-22F™ (Promega) were generated by transfecting cells with Lipofectamine 3000 following the manufacturer’s instructions. 24 h post-transfection, the antibiotic Hygromycin B (50 µg/mL; Sigma-Aldrich) was added to culture medium to select for successful GloSensor-22F-integrated GLUTag cells. Stable transfected cells were passaged several times before experimental use.

### 2.3. Expression analysis

RNA sequencing of FACS-purified, Venus-expressing cells from the colon of GLU-Venus mice as described previously [26], [27]. All sequencing was conducted at the Transcriptomics and Genomics Core Facility (Cancer Research UK Cambridge Institute) using an Ilumina HiSeq 2500 system.

### 2.4. GLP-1 secretion assay

Primary colonic cultures or GLUTag cells were plated at a density of 160 000 cells/well on 2% BME-coated 24-well plates the night prior. When applicable, cultures were preincubated with 0.5 µg/mL pertussis toxin (Cayman Chemical) for 4 h on the day of the experiment. Following incubation, cells were washed three times with warmed standard saline (138 buffer supplemented with 0.1 mM D-glucose and 0.1% BSA) for 30 min at 37°C. Standard saline solution was removed, and application of drugs were incubated for 2 h at 37°C and secretion supernatants were collected and centrifuged at 2000 RPM for 5 min at 4°C. The resulting supernatant was transferred to a new tube and snap frozen prior to analysis. Remaining cells were lysed and centrifuged at 10 000 RPM for 10 min at 4°C, transferred to a new tube and snap frozen prior to analysis. Total GLP-1 was quantified by ELISA (BMS2194, Invitrogen) following the manufacturer’s instructions and measured on the FlexStation 3 (Molecular Devices) plate reader. In GLUTag cells, GLP-1 was normalized to total protein measured from cell lysates and expressed as a percentage of average basal secretion or forskolin secretion for each experiment, as indicated in the Figure legends. For primary secretion experiments, GLP-1 secretion was calculated as a percentage of total hormone content per well.

### 2.5. cAMP transfection and live-cell cAMP measurement

GLUTag cells stably expressing the cAMP biosensor GloSensor-22F™ (Promega) were generated for experimental use. Prior to experiments, cells were plated at a density of 80 000 cells/well on 2% Matrigel-coated black, clear-bottom 96-well plates (ThermoFisher) for 16-18 h. Using a modified protocol [28], cells were washed three times with warmed standard saline (138 buffer supplemented with 0.1 mM D-glucose and 0.1% BSA) and loaded with 80 µL/well D-luciferin (2mM; Gold Biotechnology) for 2 h at room temperature (20-23°C) in the dark. Luminescence measurements were obtained using the Tecan Spark^TM^ Cyto plate reader at 20.5-21.5°C, integration 1.0 s, and interval 1.5 min. Luminescence intensity was measured after the addition of select treatments and following the addition of 10 µM forskolin. A 3-point rolling ball average filter was applied at the end of each treatment and after the addition of 10 µM forskolin. cAMP production was determined at the end of each treatment or after the addition of 10 µM forskolin by normalizing luminescence intensity measured to average vehicle control for each experiment. Rate of cAMP production was determined by calculating the slope at the end of each treatment or after the addition of 10 µM forskolin by normalizing luminescence measured to average vehicle control for each experiment.

### 2.6. Ca^2+^ imaging

Cells were loaded with 5 µM Fura-2-acetoxymethyl ester (Fura-2AM, Invitrogen) for 40 min in standard saline (138 buffer supplemented with 0.1 mM D-glucose). Cells were washed seven times with warmed standard saline before Fura2 calcium imaging was performed on a PTI system connected to a Nikon TE2000-U Eclipse inverted microscope equipped with a 40X water objective (1.2 NA). Fluorescence measurements following sequential excitation at 340 nm and 380 nm were acquired with the PCO-edge mono CMOS camera (Excelitas) every 2.5 s using EasyRatio Pro software (version 3.4.130.86, Horiba). Cells were continuously perfused during recordings at a rate of 1 mL/min with standard saline or drug using a gravity-fed perfusion system. Experiments were performed at room temperature (20-24 °C). The Fura 2 ratio (340/380) was calculated from the fluorescence intensity measurements in a user-selected area marked inside the cell that was background corrected. A four-point rolling ball average filter was applied to the data to minimize noise and smooth out fluctuations. For data inclusion, only cells with a noise level within ± 0.1, a SD of <0.05 from beginning of recording to maximal response and a mean change in fluorescence ratio (ΔFura2) >0.2 during maximal response were considered. ΔFura2 refers to the magnitude of the maximal Ca^2+^ responses during drug application minus the baseline average signal 20s prior drug application. The threshold for Ca^2+^ responses was set as ΔFura2 >0.05.

### 2.7. Solutions, drugs and chemicals

Standard saline solution (138 buffer) contained (in mM): NaCl (138), KCl (4.5), HEPES (10.0), NaHCO_3_ (4.2), NaH_2_PO_4_ (1.2), CaCl_2_ (2.6), MgCl_2_ (1.2); pH 7.4 with NaOH. Secretion and cAMP imaging experiments were supplemented with 0.1 mM D-glucose and 0.1% BSA, and Ca^2+^ imaging experiments were supplemented with 0.1 mM D-glucose.

Unless otherwise stated, all chemicals were purchased from MilliporeSigma. 4-CMTB, somatostatin and Y-27632 was purchased from Tocris Bioscience. YM-254890 was purchased from Cayman Chemical. Drug stocks were made as 1000× concentrated stock and were diluted to a final working concentration on the day of the experiment.

### 2.8. Data Analysis

Details of specific statistical analysis testing and outputs are detailed in corresponding Figure legends. Statistical tests were performed with GraphPad Prism 10 (GraphPad Software). Statistical significance between two groups was determined using the Wilcoxon signed rank test, as indicated. Statistical significance to compare three or more groups was determined using either a parametric one-way analysis of variance (ANOVA) test followed by the Dunnett’s or Sidak’s test for multiple comparisons or a non-parametric Kruskal-Wallis test followed by the Dunn’s test for multiple comparisons. *P <* 0.05 was considered statistically significant.

## 3. Results

### 3.1. SCFAs differentially modulate the release of GLP-1 from EECs

Prior to investigating the signaling mechanisms recruited by the free fatty acid receptor 2 and 3 (FFA2 and FFA3) in GLP-1 releasing EECs, we first analyzed the expression profile of several known SCFA-sensitive receptors in cells. Using transgenic mice with fluorescently labelled GLP-1 releasing cells [29], GLP-1 releasing cells were isolated and enriched using fluorescence-activated cell sorting (FACS) before processing for bulk RNA sequencing. Given the abundance of SCFAs within the distal gut where gut bacteria reside, we chose to focus on physiologically relevant colonic GLP-1 releasing cells in our analysis. Expression of *Ffar2* and *Ffar3* were confirmed in colonic mouse GLP-1-releasing EECs at levels comparable to the well-characterized bombesin receptor 2 (encoded by *Grpr*) [30] (Figure 1A) [31]. We also sought to examine the expression profile of SCFA-sensitive receptors in a cell line model of colonic GLP-1 releasing EECs, the GLUTag cell line [32]. *Ffar2* and *Ffar3* expression in the GLUTag cell line closely mirrored that of primary cells (Figure 1B). Other SCFA-sensitive receptors showed negligible expression in GLUTag cells, thus supporting the use of GLUTag cells as a model for studying specific FFA2 and FFA3 signaling mechanisms in GLP-1-releasing EECs.

**Figure 1:**
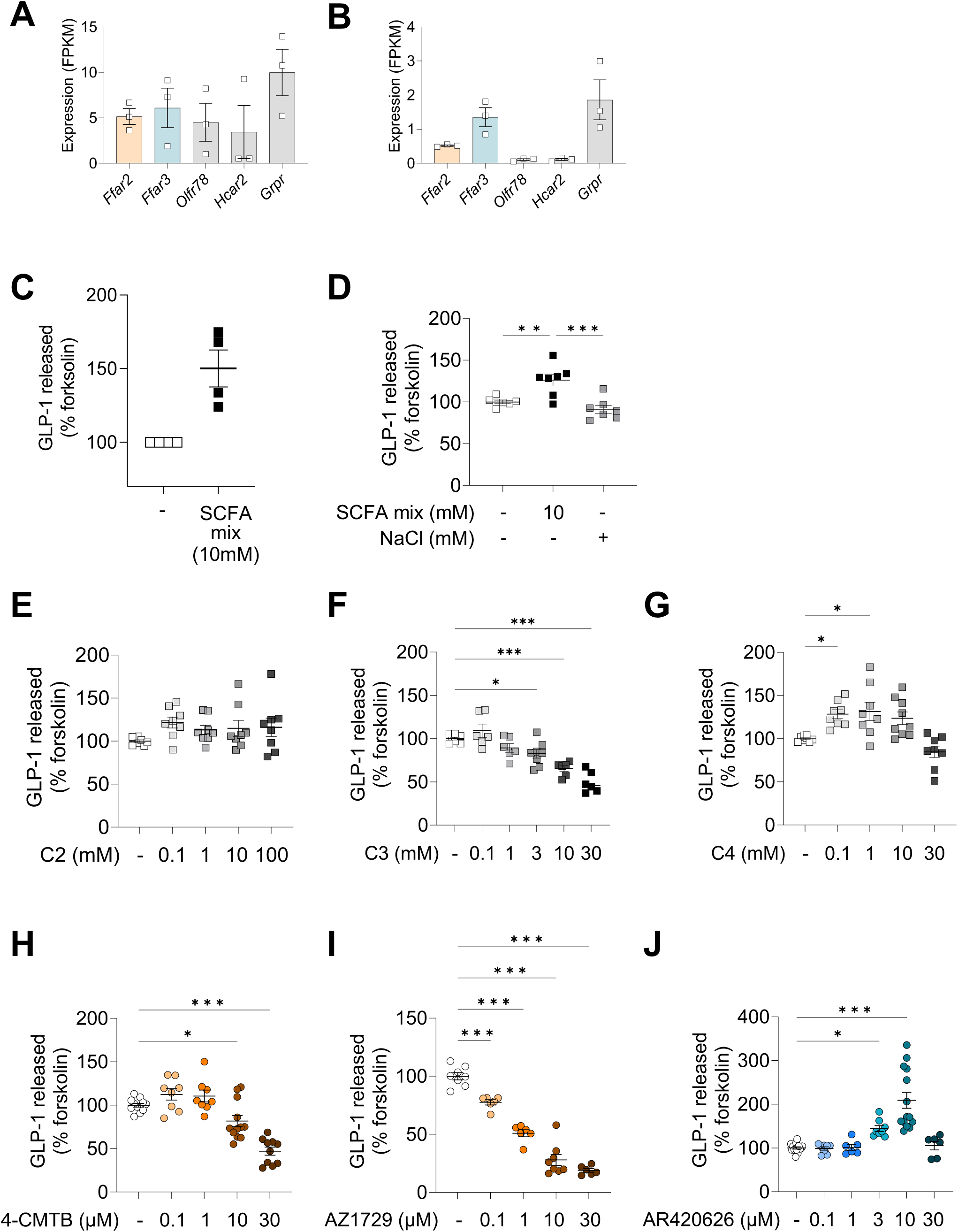
Expression of SCFA-sensitive GPCRs in GLP-1 releasing EEC and GLP-1 secretion responses. Expression analysis of SCFA-sensitive receptors in a FACS-enriched population of colonic mouse GLP-1 releasing EEC (A) or GLUTag cells (B). Individual symbols represent transcripts per million (TPM) from 10,000 FACS enriched cells per biological replicate. Mean ± SEM indicated. GLP-1 secretion levels from primary mouse colonic cultures (C) or (D) GLUTag cells following 2h treatment with a mild stimulus (2 µM forskolin) and the addition a SCFA mixture (10 mM). GLP-1 levels released from GLUTag cells following 2h treatment with a mild stimulus (2 µM forskolin) and the addition of individual SCFAs including acetate (E), propionate (F) or butyrate (G). GLP-1 secretion levels from GLUTag cells following 2h treatment with a mild stimulus (2-5 µM forskolin) and a selective ligand of FFA2, 4-CMTB (H) or AZ1729 (I), or FFA3, AR420626 (J). For primary secretion experiments, GLP-1 secretion was calculated as a percentage of total hormone content per well. All GLUTag cell secretion data was normalized to total protein from cell lysates and expressed relative to the forskolin alone condition. Individual symbols represent GLP-1 released per well. Orange symbols represent responses targeting FFA2, blue symbols represent responses targeting FFA3, grey symbols represent potential activation of multiple SCFA-sensitive receptors. N = 6-18 from at least 3 independent experiments, mean ± SEM indicated. One-way ANOVA with Dunnett’s multiple comparisons statistical test performed; *\*P<0.05, **P<0.01, ***P<0.001*.

Having confirmed receptor expression, we next examined the functional effects of SCFA treatment on GLP-1 release in primary mouse colonic cultures and GLUTag cells. In the presence of a mild stimulus (forskolin 2-5 µM), a SCFA mix significantly increased GLP-1 secretion in both colonic preparations (Figure 1C+D). To identify which SCFAs contributed to this effect, we tested acetate (C2), propionate (C3) and butyrate (C4) alone on GLUTag cells. In the absence of forskolin, each SCFA produced only mild effects on GLP-1 release (Supplementary Figure 1A-C). However, with the addition of forskolin, distinct SCFA-induced effects emerged (Figure 1E-G). Acetate did not significantly affect GLP-1 release, even in the presence of forskolin (Figure 1E). Propionate, in contrast, significantly reduced GLP-1 release in a dose-dependent manner (Figure 1F), whereas butyrate induced a biphasic response, enhancing secretion at lower concentrations, but having no effect at higher concentrations (Figure 1G). These findings suggest SCFAs induce receptor-specific effects, with the overall effect observed on GLP-1 secretion likely driven by differences in their agonist potency profiles.

To further investigate receptor-specific signaling mechanisms, we selectively activated FFA2 or FFA3 with pharmacologically selective ligands. We tested the selective FFA2 ligands 4-CMTB and AZ1729, the latter of which is reported to bias downstream Ga_i_-signaling pathways [33]. Increasing concentrations of 4-CMTB alone did not significantly alter GLP-1 secretion (Supplementary Figure 1D). However, in the presence of forskolin, 4-CMTB treatment markedly reduced GLP-1 release in a dose-dependent manner (Figure 1H). Similarly, AZ1729 significantly inhibited GLP-1 secretion under both conditions: treatment of AZ1729 alone (Supplementary Figure 1E) and treatment of AZ1729 in the presence of forskolin (Figure 1I). These findings suggest that FFA2 activation inhibits GLP-1 release. Conversely, selective activation of FFA3 using AR420626 dose-dependently increased GLP-1 release up to 10 µM, with no further increases at higher concentrations (Supplementary Figure 1F). This response was consistent in the presence of forskolin (Figure 1J), supporting FFA3 activation as a stimulator of GLP-1 release. To better understand the differential effects of FFA2 and FFA3 on GLP-1 release, we investigated the downstream signaling pathways recruited by each selective ligand in GLP-1-releasing EECs.

### 3.2. FFA2 activation by 4-CMTB increases cAMP production independent of Ga_i-_ and Ga_q_-protein coupling in GLP-1 releasing EECs

To investigate the inhibition of GLP-1 release observed in GLUTag cells following treatment with 4-CMTB, we first examined the possible involvement of intracellular cAMP. This second messenger is involved in regulating GLP-1 release from EECs, and its production is inhibited by Ga_i_-proteins, to which the receptor FFA2 has been shown to couple to in overexpression studies [20]. Using transfected GLUTag cells stably expressing the GloSensor 22F™ cAMP biosensor [34], we recorded the luminescence signal generated from live GLUTag cells treated with different concentrations of 4-CMTB. Forskolin, a direct activator of adenylyl cyclase was added after to elevate cAMP levels. An independent group of GloSensor-expressing GLUTag cells was treated with somatostatin (SST), which activates somatostatin receptors expressed on GLP-1 releasing EECs that exclusively couple to Ga_i_-proteins, serving as a positive control for Ga_i_-coupling. After application of 4-CMTB, there was no change in cAMP levels measured except at the highest concentration tested (30 µM, Supplementary Figure 2A). However, during forskolin stimulation a concentration-dependent increase in cAMP levels was observed in 4-CMTB treated groups (Figure 2A+B), which was significant compared to the vehicle control group. Similar results were observed when examining rates of cAMP production following application of 4-CMTB or forskolin (Supplementary Figure 2B+C).

**Figure 2:**
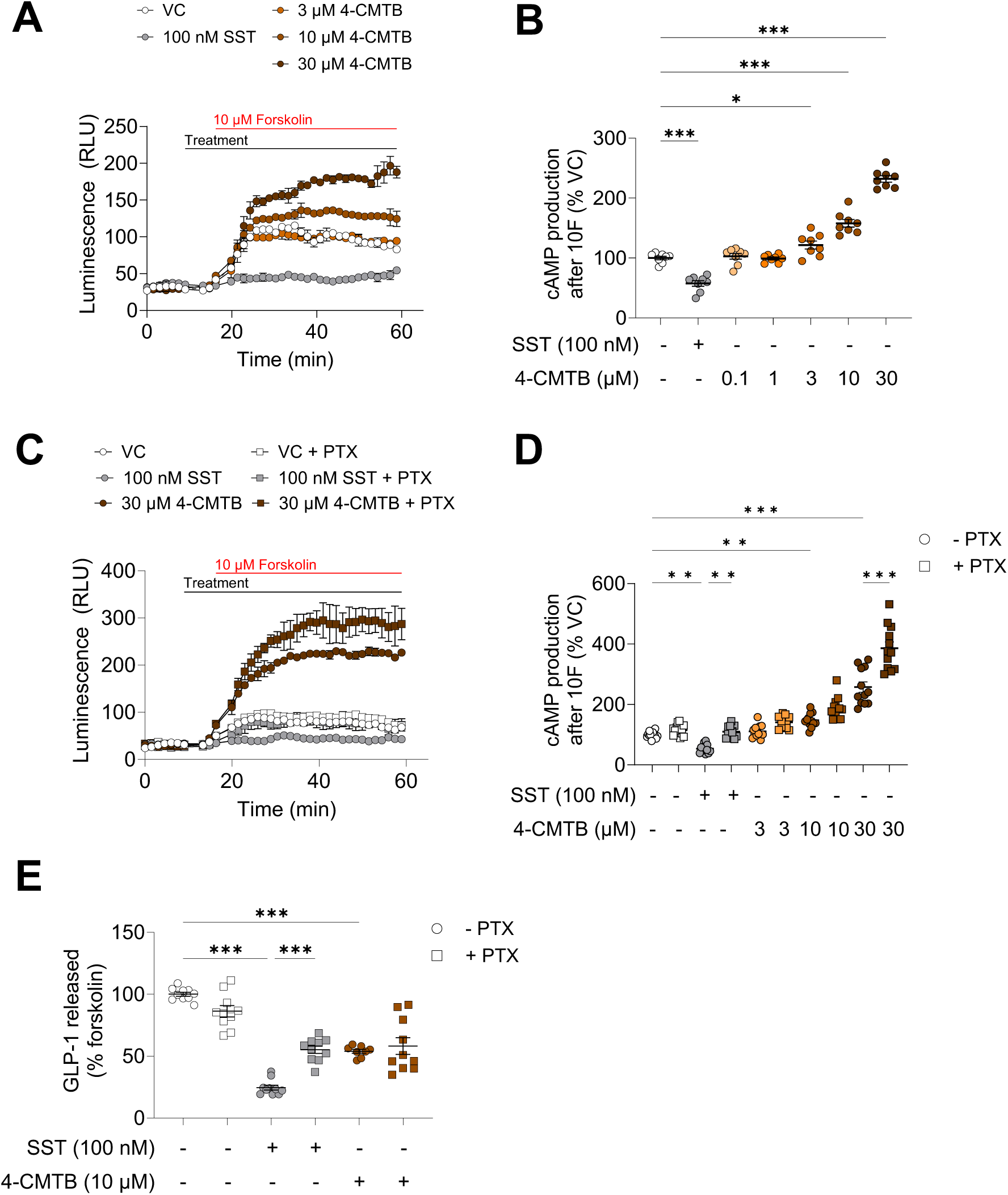
4-CMTB increases cAMP levels in GLUTag cells independent of Ga_i_-protein coupling. Representative traces of live-cell cAMP measurements following 4-CMTB treatment in the absence (A) or presence of pertussis toxin (PTX; 0.5 µg/mL for 4h) (B). Intracellular cAMP levels monitored as luminescence (relative luminescence units, RLU) over time in GLUTag cells stably-expressing GloSensor-22F™. Addition of treatments and a mild stimulus (10 µM forskolin) shown in black and red labelled bars, respectively. Calculated peak cAMP production, determined as the maximal response of each treatment after the addition of forskolin normalized to vehicle control, in response to 4-CMTB treatments in the absence (C) or presence of PTX (D). Individual symbols represent a single well, PTX-treated wells indicated by square symbols. N = 8-14 from at least 3 independent experiments, mean ± SEM indicated. One-way ANOVA with Dunnett’s (C) or Sidak’s (D) multiple comparisons statistical test performed; *\*P<0.05, **P<0.01, ***P<0.001*. GLP-1 levels released from GLUTag cells following 2h treatment with a mild stimulus (2 µM forskolin) and 4-CMTB in the absence or presence of PTX (E). All secretion data was normalized to total protein from cell lysates and expressed relative to the forskolin alone condition. Individual symbols represent GLP-1 released per well. N = 8-10 from at least 3 independent experiments, mean ± SEM indicated. One-way ANOVA with Sidak’s multiple comparisons statistical test performed; *\*P<0.05, **P<0.01, ***P<0.001*.

To delineate the G-protein involved in 4-CMTB stimulated cAMP accumulation, we utilized the Ga_i_-uncoupler, pertussis toxin (PTX; 0.5 µg/mL pre-treatment for 4h). Pre-treatment with PTX did not affect cAMP accumulation in the vehicle control treatment group, but amplified cAMP production induced by 4-CMTB treatment at the higher concentrations tested (Figure 2C+D), suggesting some recruitment of Ga_i-_protein-coupled mechanisms downstream of FFA2 activation. However, the lack of effect of PTX pre-treatment on GLP-1 secretion from GLUTag cells (Figure 2E) suggests minimal involvement of cAMP production on reduced GLP-1 release from GLUTag cells following FFA2 activation by 4-CMTB.

The involvement of Ga_q_, the other G protein family shown to couple to FFA2 [20], was assessed using the Ga_q_-blocker YM-254890 (10 pM). We found cAMP production following 4-CMTB treatment was unchanged in the presence of the Ga_q_-blocker YM-254890 (Figure 3A+B). Furthermore, YM-254890 did not block the inhibition of GLP-1 release by 4-CMTB but significantly reduced the stimulatory effect of the Ga_q_-coupled receptor agonist, bombesin (BBS; Figure 3C), demonstrating the selectivity of YM-254890 to block downstream Ga_q_-signaling. These results suggest that the increased cAMP production by 4-CMTB does not involve Ga_q_ signaling mechanisms.

**Figure 3:**
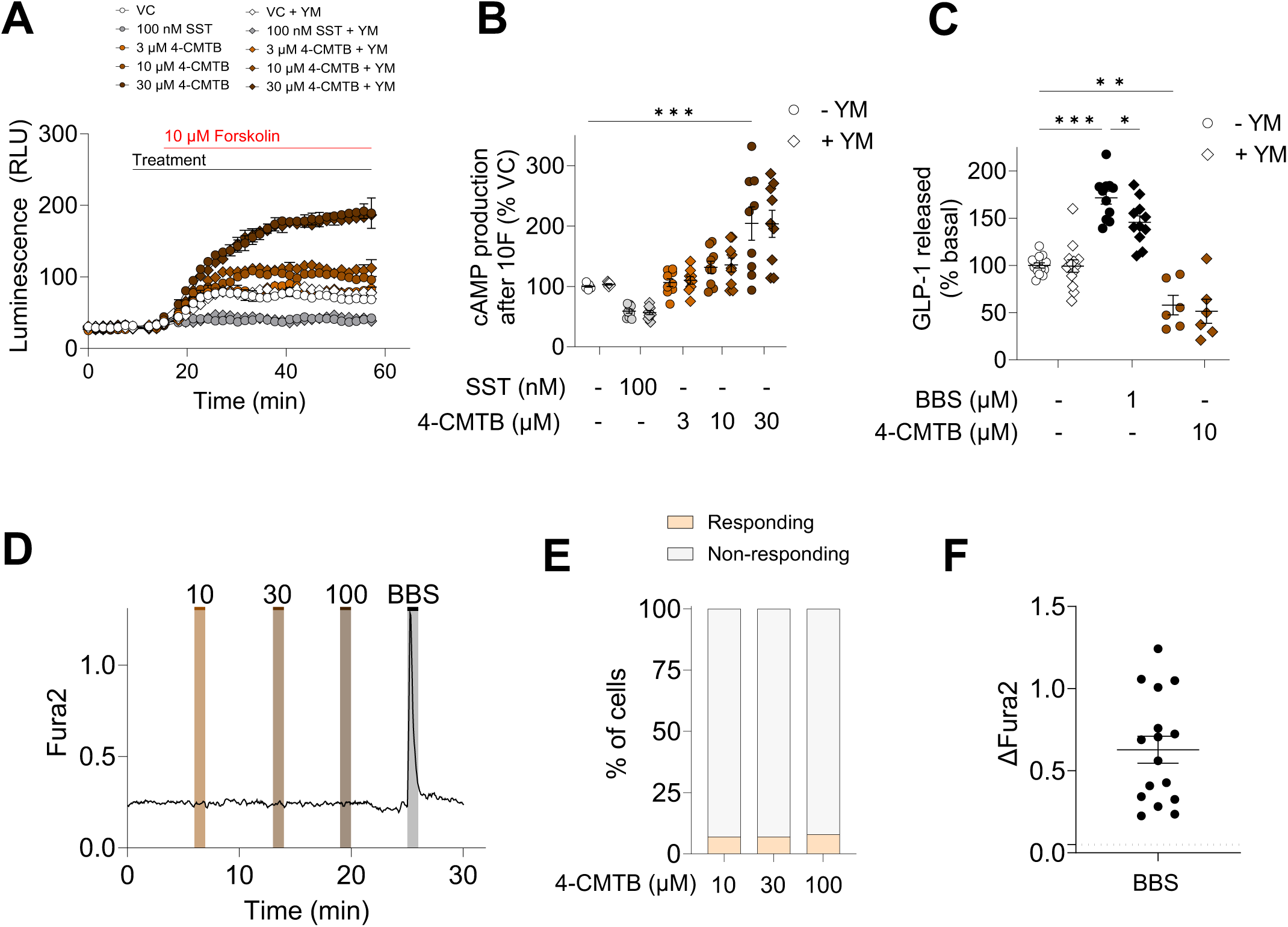
4-CMTB increases cAMP levels in GLUTag cells independent of Ga_q_-protein coupling and does not alter intracellular Ca^2+^ levels. Representative traces of live-cell cAMP measurements following 4-CMTB treatment in the absence or presence of YM-254890 (YM; 10 pM) (A). Similar experimental design and analysis performed as detailed in Figure 2. Calculated peak cAMP production in response to 4-CMTB in the absence and presence of YM (B). Individual symbols represent a single well, YM-treated wells indicated by diamond symbols. N = 8-9 from at least 3 independent experiments, mean ± SEM indicated. One-way ANOVA with Sidak’s multiple comparisons statistical test performed; *\*P<0.05, **P<0.01, ***P<0.001*. GLP-1 levels released from GLUTag cells following 2h treatment with a mild stimulus (2 µM forskolin) and the addition of the 4-CMTB in the absence or presence of YM (C). All secretion data was normalized to total protein from cell lysates and expressed relative to the forskolin alone condition. Individual symbols represent GLP-1 released per well. N = 6-13 from at least 3 independent experiments, mean ± SEM indicated. One-way ANOVA with Sidak’s multiple comparisons statistical test performed; *\*P<0.05, **P<0.01, ***P<0.001*. Representative recording of intracellular Ca^2+^ levels during treatment with 4-CMTB (10 - 100 µM) and 100 nM bombesin (BBS) (D). Intracellular Ca^2+^ measured using ratiometric Fura2 fluorescence at 340 and 380 nm excitation. Drugs were applied as indicated above trace. Calculated % of GLUTag cells responding to 4-CMTB treatment with an increase in Fura2 fluorescence above a pre-set threshold (>0.05) (E). Orange bar indicates % of cells that responded to 4-CMTB treatment, grey bar indicates % of cells that did not respond to 4-CMTB treatment. N = 8-16 from at least 3 independent experiments. Change in Fura2 fluorescence ratio in response to 100 nM BBS (F). Individual data points represent individual GLUTag cells, lines represent mean ± SEM. Dotted line represents Fura2 threshold (0.05). N = 16 from at least 3 independent experiments.

### 3.3. FFA2 activation by 4-CMTB does not trigger intracellular Ca^2+^ release in GLP-1 releasing EECs

To further investigate possible involvement of Ga_q_-protein signaling pathways in GLP-1 releasing EECs, the canonical second messenger signaling pathway associated with Ga_q_-protein activated pathways, intracellular Ca^2+^ released from ER stores, was assessed by Fura2 imaging. In GLUTag cells, increasing concentrations of 4-CMTB failed to elicit detectable changes in intracellular Ca^2+^ levels (Figure 3D+E). While 4-CMTB did not induce Ca^2+^ responses, robust Ca^2+^ responses were detected upon BBS treatment (Figure 3F). Combined with the negligible effect of YM-254890 on 4-CMTB inhibited release of GLP-1, activation of FFA2 by 4-CMTB does not recruit canonical Ga_q_ protein signaling mechanisms.

### 3.4. FFA2 activation by AZ1729 decreases cAMP production independent of Ga_i_- and Ga_z_-protein coupling in GLP-1 releasing EECs

The FFA2 allosteric agonist AZ1729, has been shown to preferentially recruit Ga_i_-signaling pathways [33]. Given the involvement of cAMP in regulating exocytosis and AZ1729 inhibition of GLP-1 release, we expected cAMP levels in GLP-1-releasing EECs would be reduced following AZ1729 treatment. Using the same cAMP detection system described previously, we assessed cAMP signaling dynamics following AZ1729 treatment in live GLUTag cells stably expressing GloSensor 22F™. Overall, a biphasic response was observed: after application of AZ1729 treatment, at lower AZ1729 concentrations (<1 µM) the rate of cAMP production significantly increased but at higher concentrations (>10 µM) the peak and rate of cAMP production significantly decreased (Figure 4A, Supplementary Figure 2D-E). This biphasic response was more pronounced during forskolin stimulation, where peak cAMP levels and rate of cAMP production were significantly decreased compared to vehicle control at lower AZ1729 concentrations and significantly increased at higher AZ1729 concentrations (Figure 4B, Supplementary Figure 2F).

**Figure 4:**
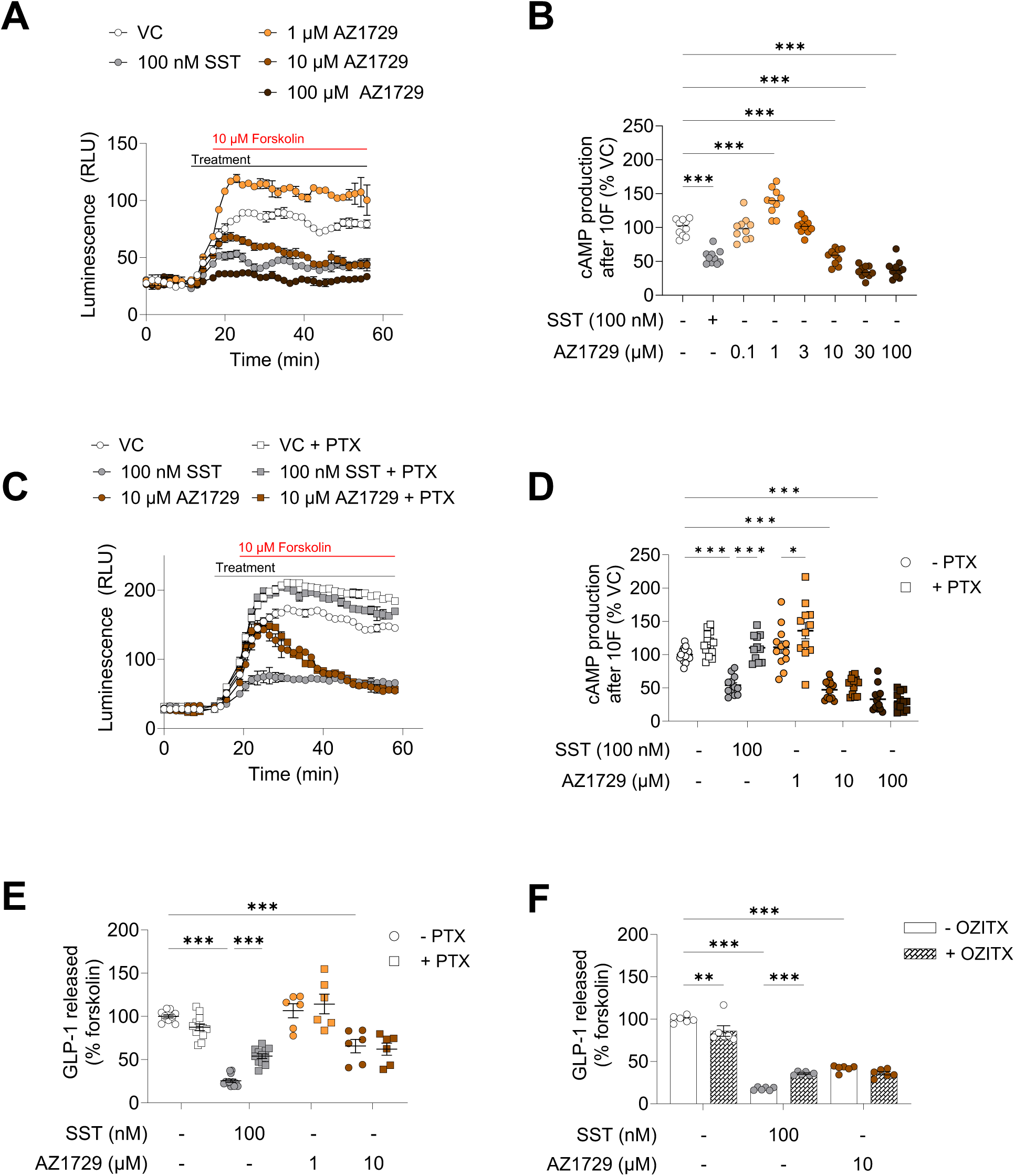
AZ1729 decreases cAMP production and GLP-1 release independent of Ga_i/o_ and Ga_z_ coupling. Representative traces of live-cell cAMP measurements following AZ1729 treatment in the absence (A) or presence of pertussis toxin (PTX; 0.5 µg/mL for 4h) (B). Similar experimental design and analysis performed as detailed in Figure 2. Calculated peak cAMP production in response to AZ1729 in the absence (C) or presence of PTX (D). Individual symbols represent a single well, PTX-treated wells indicated by square symbols. N = 10-16 from at least 3 independent experiments, mean ± SEM indicated. One-way ANOVA with Dunnett’s (C) or Sidak’s (D) multiple comparisons statistical test performed; *\*P<0.05, **P<0.01, ***P<0.001*. GLP-1 levels released from GLUTag cells following 2h treatment with a mild stimulus (2 µM forskolin) and AZ1729 in the absence or presence of PTX (E) or OZITX (F). All secretion data was normalized to total protein from cell lysates and expressed relative to the forskolin alone condition. Individual symbols represent GLP-1 released per well. N = 6-12 from at least 3 independent experiments, mean ± SEM indicated. One-way ANOVA with Sidak’s multiple comparisons statistical test performed; *\*P<0.05, **P<0.01, ***P<0.001*.

To determine the involvement of Ga_i_-protein coupling in mediating AZ1729 responses, we treated GLUTag cells with PTX during AZ1729 application and assessed the effects on cAMP production and GLP-1 release. Interestingly, AZ1729-mediated inhibition of cAMP production and GLP-1 release persisted in the presence of PTX (Figures 4C+D). PTX treatment was able to attenuate SST-inhibition of cAMP production and GLP-1 release (Figure 4C-E), confirming functional Ga_i_- coupling in GLUTag cells. Possible involvement of PTX-insensitive Ga_z_-protein coupling in AZ1729 responses was also assessed using the broad Ga_i/o/z_-uncoupler, OZITX [35]. OZITX expression attenuated SST-induced inhibition of GLP-1 release but had no effect on AZ1729-mediated inhibition of GLP-1 release in GLUTag cells (Figure 4F). This supports our findings that FFA2-activation by AZ1729 decreased cAMP production and GLP-1 release by a novel, non-Ga_i_-protein family coupling mechanism.

Intracellular Ca^2+^ signaling mechanisms may regulate cAMP levels [36], so the effectiveness of AZ1729 in triggering Ca^2+^ signaling mechanisms was investigated using Fura2 Ca^2+^ imaging as described previously. We found increasing concentrations of AZ1729 failed to elicit detectable changes in intracellular Ca^2+^ levels in GLUTag cells (Supplementary Figure 3). Therefore, the effect of AZ1729 on intracellular cAMP levels could not be attributed to intracellular Ca^2+^ signaling mechanisms.

### 3.5. FFA3 activation by AR420626 does not alter cAMP production in GLP-1 releasing EECs

Following our investigation of FFA2-selective ligands, we explored FFA3 activation using the selective agonist AR420626. Activation of FFA3 by AR420626 stimulated GLP-1 release in GLUTag cells (Figure 1H), prompting us to investigate cAMP production in live GLUTag cells stably expressing GloSensor22F™. Increasing AR420626 concentrations did not enhance peak or rates of cAMP production compared to the vehicle control (Supplementary Figure 2G+H) and no change was observed in peak or rate of cAMP production during forskolin stimulation compared to vehicle control (Figure 5A+B, Supplementary Figure 2I). PTX treatment was unable to unmask an effect of increasing concentrations of AR420626 on cAMP production (Figure 5C+D). Although, AR420626 stimulation of GLP-1 release was attenuated by PTX treatment (Figure 5E), this suggests Ga_i_-signaling involvement in GLP-1 secretion independent of cAMP production.

**Figure 5:**
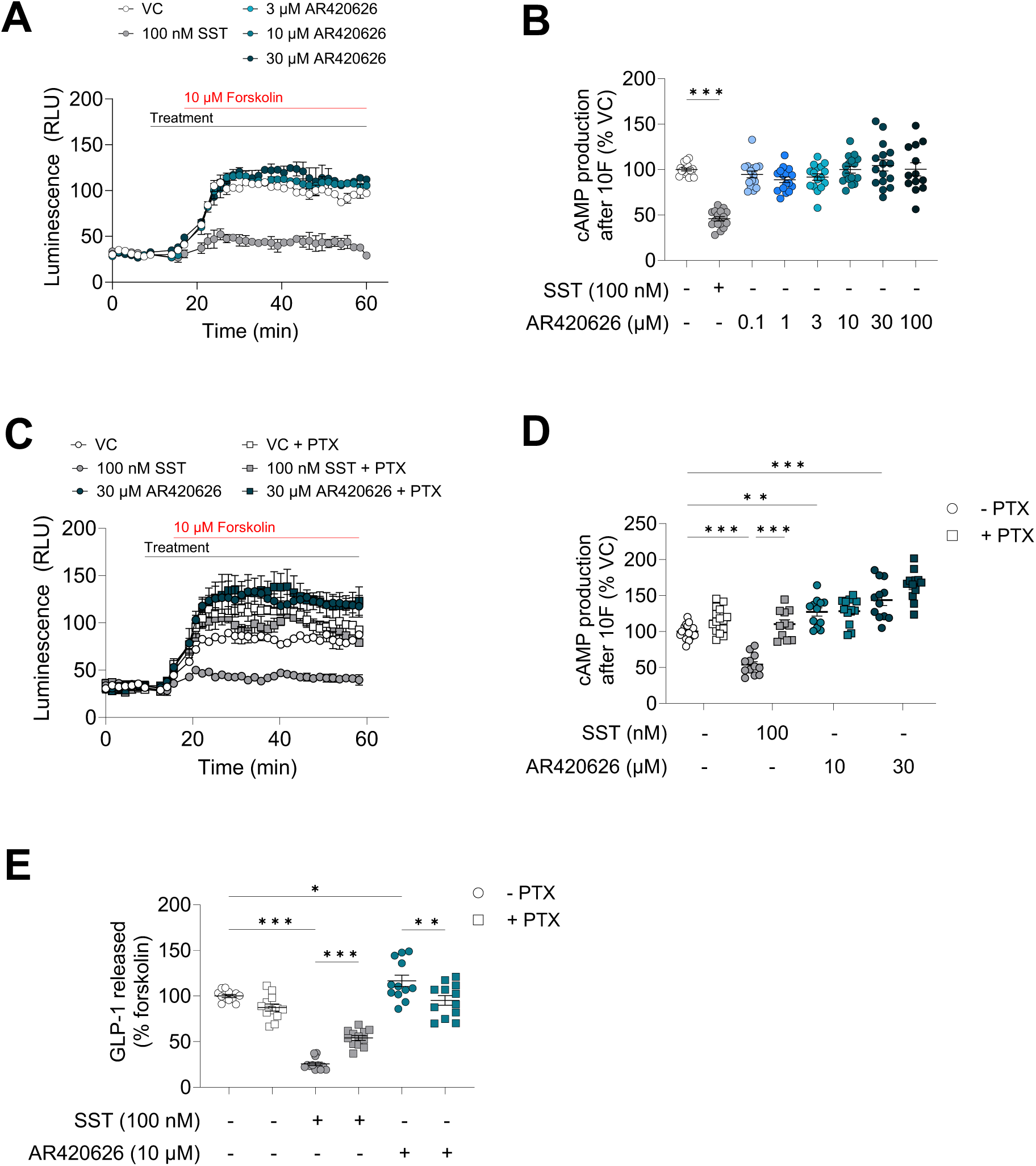
Blockade of Ga_i_-protein coupling reveals AR420626 induced increase in cAMP levels and GLP-1 release from GLUTag cells. Representative traces of live-cell cAMP measurements following AR420626 treatment in the absence (A) or presence of pertussis toxin (PTX; 0.5 µg/mL for 4h) (B). Similar experimental design and analysis performed as detailed in Figure 2. Calculated cAMP production in response to AR420626 in the absence (C) or presence of PTX (D). Individual symbols represent a single well, PTX-treated wells indicated by square symbols. N = 10-16 from at least 3 independent experiments, mean ± SEM indicated. One-way ANOVA with Dunnett’s (C) or Sidak’s (D) multiple comparisons statistical test performed; *\*P<0.05, **P<0.01, ***P<0.001*. GLP-1 levels released from GLUTag cells following 2h treatment with a mild stimulus (2 µM forskolin) and AR420626 in the absence or presence of PTX (E). All secretion data was normalized to total protein from cell lysates and expressed relative to the forskolin alone condition. Individual symbols represent GLP-1 released per well. N = 12 from at least 3 independent experiments, mean ± SEM indicated. One-way ANOVA with Sidak’s multiple comparisons statistical test performed; *\*P<0.05, **P<0.01, ***P<0.001*.

### 3.6. AR420626-induced FFA3 activation triggers an increase in intracellular Ca^2+^ levels in GLP-1 releasing EECs

To address which second messenger signaling pathways contributed to GLP-1 release following FFA3 activation, we measured intracellular Ca^2+^ responses to AR420626 application using the calcium-sensitive dye Fura2. AR420626 induced modest, concentration-dependent increases in intracellular Ca^2+^ levels (Figure 6A), with detectable responses at 10 µM (10/16 cells, mean

**Figure 6:**
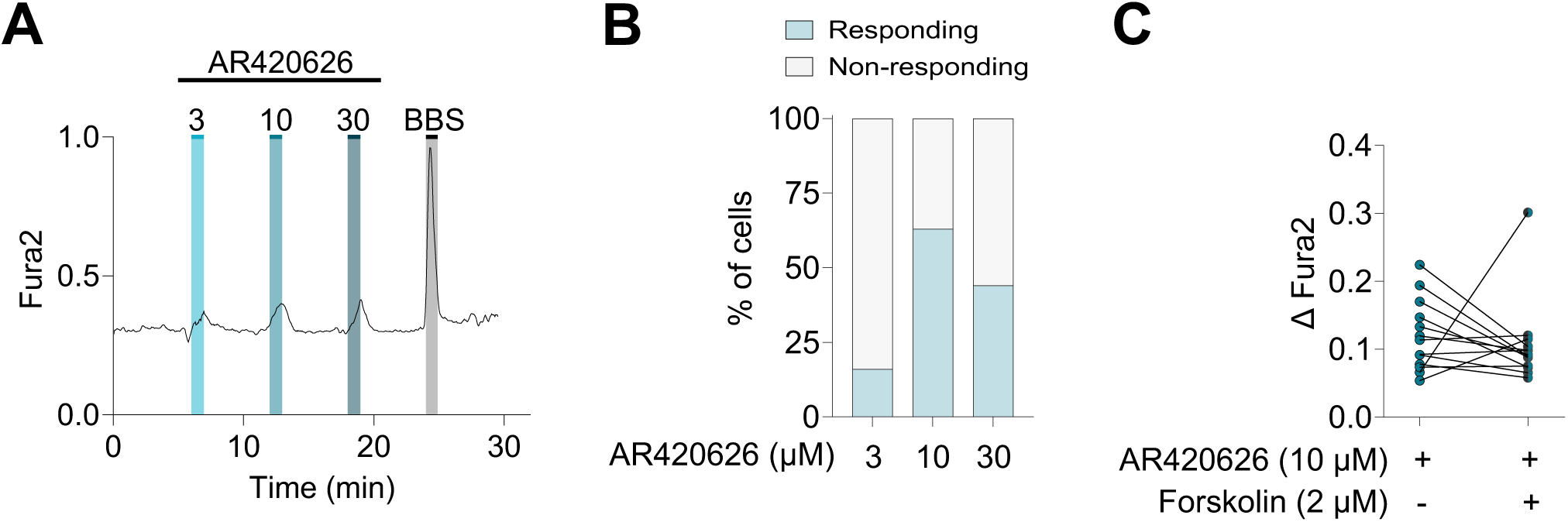
AR420626 increases intracellular Ca^2+^ levels in GLUTag cells. Representative recording of intracellular Ca^2+^ levels during treatment with AR420626 (3 - 30 µM) and 100 nM bombesin (BBS)(A). Intracellular Ca^2+^ measured using ratiometric Fura2 fluorescence at 340 and 380 nm excitation. Drugs were applied as indicated above trace. Calculated % of GLUTag cells responding to AR420626 treatment with an increase in Fura2 fluorescence above a pre-set threshold (>0.05) (E). Blue bar indicates % of cells that responded to AR420626 treatment, grey bar indicates % of cells that did not respond to AR420626 treatment. N = 8-16 from at least 3 independent experiments. Change in Fura2 fluorescence ratio in response to 10 µM AR420626 with or without 2 µM Forskolin (C). Individual data points represent individual GLUTag cells, blue symbols indicates AR420626 treatment, black and blue symbols represent the co-application of AR420626 and forskolin treatments. Solid line connects paired measurements made from the same GLUTag cell. Dotted line represents Fura2 threshold (0.05). N = 16 from at least 3 independent experiments. Wilcoxon signed-rank statistical test performed.

ΔFura2 ± SEM = 0.057 ± 0.004) and 30 µM (7/16 cells, mean ΔFura2 ± SEM = 0.069 ± 0.022; Figure 6B). To assess whether these modest Ca^2+^ increases could be potentiated by forskolin, parallel experiments were performed. Forskolin, which does not independently trigger intracellular Ca^2+^ responses in GLUTag cells (mean ΔFura2 ± SEM = 0.019 ± 0.005, n = 16), failed to enhance AR420626-induced Ca^2+^ responses compared to AR420626 treatment alone (Figure 6C), suggesting that AR420626 is responsible for mediating the Ca^2+^ response in GLP-1 releasing EECs.

### 3.7. Ketone bodies differentially modulate the release of GLP-1 in GLP-1 releasing EECs

Given the overlapping roles of SCFAs and ketone bodies in metabolic regulation, we assessed their molecular similarity using the maximum common substructure Tanimoto score, which quantifies the shared structural features [37]. Acetoacetate (AcAc) and β-hydroxybutyrate (BHB) share key functional groups with SCFAs, suggesting potential ligand promiscuity between FFA2 and FFA3. We visualized structural similarity among SCFAs, MCFAs and ketone bodies in a heatmap shown in Figure 7A. SCFAs clustered together and were structurally more similar to AcAc and BHB than to MCFAs, with the exception of acetone. Therefore, further analyses focused on AcAc and BHB, supporting the potential for shared receptor interactions between SCFAs and ketone bodies.

**Figure 7:**
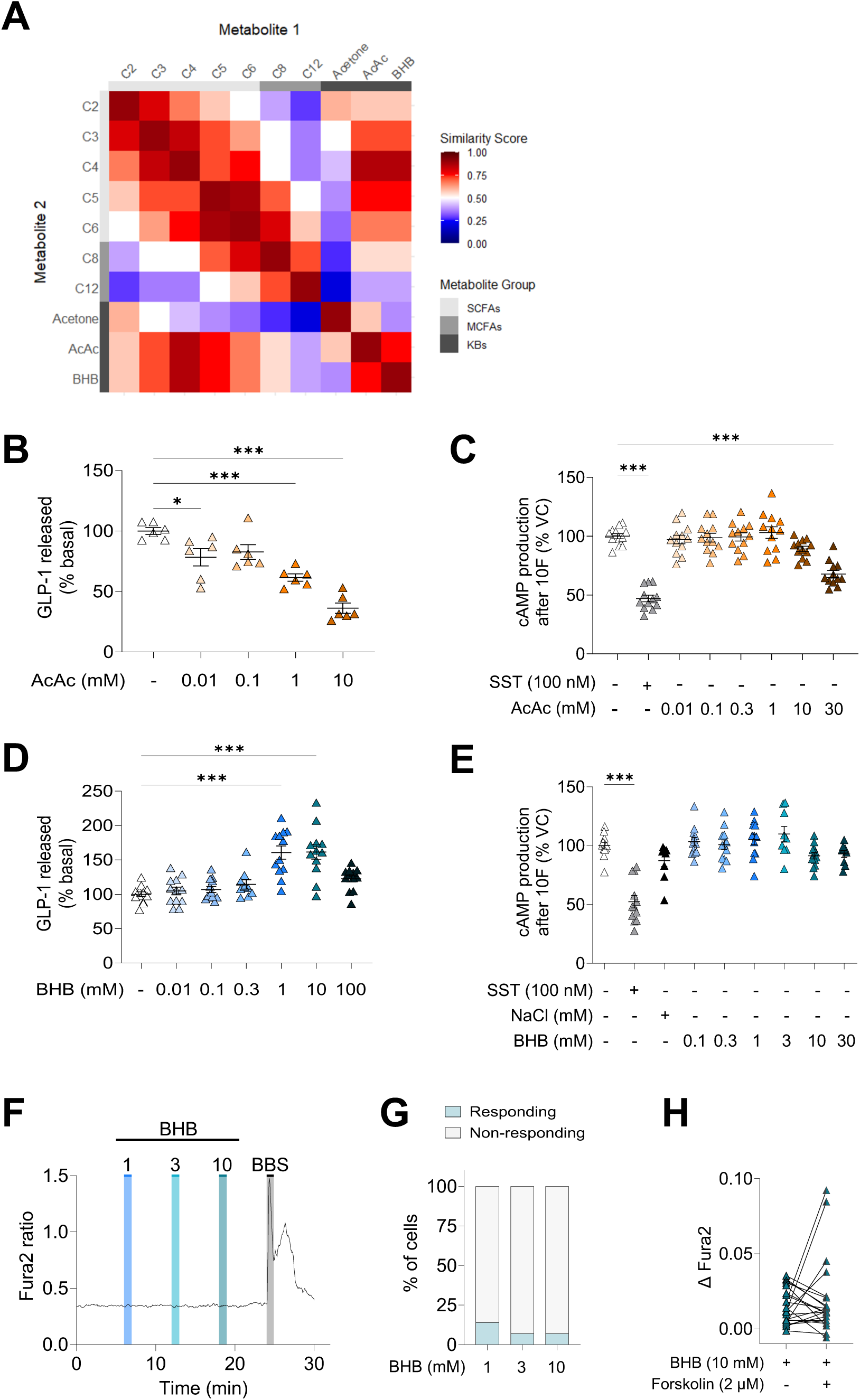
Signaling mechanisms of ketone bodies in GLUTag cells. Heatmap representation of the maximum common substructure Tanimoto scores between short-chain fatty acids (SCFAs), medium chain fatty acids (MCFAs) and ketone bodies (KB) (A). GLP-1 levels released from GLUTag cells following 2h treatment with acetoacetate (AcAc)(B) or β-hydroxybutyrate (BHB) (C). All secretion data was normalized to total protein from cell lysates and expressed relative to the standard saline condition. Individual symbols represent GLP-1 released per well. N = 6-10 from at least 3 independent experiments, mean ± SEM indicated. One-way ANOVA with Dunnett’s multiple comparisons statistical test performed; *\*P<0.05, ***P<0.001.* Calculated peak cAMP production, determined as the maximal response of each treatment after the addition of forskolin normalized to vehicle control, in response to AcAc (D) or BHB treatments (E). Individual symbols represent a single well. N = 9-12 from at least 3 independent experiments, mean ± SEM indicated. One-way ANOVA with Dunnett’s multiple comparisons statistical test performed; *\*P<0.05, **P<0.01, ***P<0.001*. Representative recording of intracellular Ca^2+^ levels during treatment with BHB (1 - 10 mM) and 100 nM bombesin (BBS) (F). Intracellular Ca^2+^ measured using ratiometric Fura2 fluorescence at 340 and 380 nm excitation. Drugs were applied as indicated above trace. Calculated % of GLUTag cells responding to BHB treatment with an increase in Fura2 fluorescence above a pre-set threshold (>0.05) (G). Blue bar indicates % of cells that responded to BHB treatment, grey bar indicates % of cells that did not respond to BHB treatment. N = 21 from at least 3 independent experiments. Change in Fura2 fluorescence ratio in response to 10 mM BHB with or without 2 µM Forskolin (H). Individual data points represent individual GLUTag cells, blue symbols indicate BHB treatment, black and blue symbols represent the co-application of BHB and forskolin treatments. Solid line connects paired measurements made from the same GLUTag cell. Dotted line represents Fura2 threshold (0.05). N = 16 from at least 3 independent experiments. Wilcoxon signed-rank statistical test performed.

To investigate ketone body interactions with FFA2 and FFA3, we examined their effects on GLP-1 secretion and second messenger signaling pathways in GLP-1-releasing EECs. AcAc dose-dependently inhibited GLP-1 release (Figure 7B) and significantly reduced peak cAMP production compared to vehicle controls during forskolin application at the highest concentration tested (Figure 7C), consistent with FFA2-mediated inhibition. However, despite this inhibitory effect, we observed an increase in cAMP levels and rate of cAMP production immediately following application of AcAc (Supplementary Figure 4A+B). BHB, in contrast, stimulated GLP-1 release at physiological concentrations (1-10 mM) (Figure 7D) but did not alter peak cAMP levels compared to vehicle controls during forskolin stimulation (Figure 7E). However, a biphasic response was observed immediately following BHB application and when determining rates of cAMP production, with higher concentrations showing some osmotic influence (Supplementary Figures 4D-F). Additionally, BHB did not trigger intracellular Ca^2+^ responses in GLUTag cells (Figures 7F+G), and responses remained unchanged in the presence of forskolin (Figure 7H). Collectively, these findings demonstrate that ketone bodies differentially modulate GLP-1 secretion via distinct signaling mechanisms.

## 4. Discussion

Nutrient-derived metabolites act as critical signaling molecules that regulate EEC function through nutrient-sensitive GPCRs [5], [26], [38], [39], [40]. Activation of these receptors controls gut hormone release, including GLP-1, which is secreted from distal small intestinal and colonic EECs to promote insulin secretion [41]. Here, we demonstrate that both SCFAs and ketone bodies signal through the SCFA receptors FFA2 and FFA3 to modulate GLP-1 secretion via distinct, non-canonical pathways.

In primary mouse colonic cultures and GLUTag cells, physiological SCFA mixtures stimulated GLP-1 secretion, supporting a role for SCFAs as activators of colonic EECs. However, responses to individual SCFAs varied: acetate had no effect, butyrate increased secretion without further dose-dependence and propionate inhibited GLP-1 release, despite previously reported stimulatory effects in mouse primary colonic cultures [5], [42] and STC-1 cells [43], while no effect was has been observed in human NCI-H716 cells [44]. Such variability may reflect basolateral localization of the SCFA receptors, which limit luminal exposure [5], [16], [45], [46], or differences between clonal cell systems and intact tissue, where paracrine interactions influence EEC activity. These results suggest that the coexistence of FFA2 and FFA3 within the same EECs, despite ∼40% sequence similarity, allows for integration of distinct nutrient cues and flexibility in secretory output, with expression levels or signaling bias potentially shifting across metabolic states. Collectively, our findings demonstrate that although individual SCFAs exert differential effects, the integrated action of SCFAs in the colon is stimulatory, underscoring their role in nutrient-dependent regulation of GLP-1 secretion.

The FFA2-mediated GLP-1 inhibition observed in our colonic EECs challenged the canonical view of the predominant Ga_q_-FFA2 coupling, which mobilizes Ca^2+^ and promotes GLP-1 release in mouse models [5], [6]. Instead, our findings demonstrate ligand-specific inhibition, likely reflecting distinct FFA2 conformations stabilized by different ligands, which could explain discrepancies with prior reports and regional variability across the intestine. The allosteric agonist 4-CMTB reduced GLP-1 secretion independent of PTX treatment yet paradoxically increased cAMP in a PTX-sensitive manner, while the Ga_i_-biased ligand AZ1729 both inhibited GLP-1 release and cAMP in a PTX-insensitive manner. The inability of OZITX, a Ga_i/o/z_-family uncoupler [35], to reverse AZ1729-mediated inhibition further supports non-canonical FFA2 coupling (Figure 4F). Consistent with these inhibitory outcomes, 4-CMTB neither triggered Ca^2+^ mobilization nor enhanced GLP-1 release in the presence of the Ga_q_ inhibitor YM-254890, supporting the absence of classical Ga_q_-coupled pathways. These findings support prior reports that demonstrated 4-CMTB fails to elicit Ca^2+^ responses in some FFA2-expressing systems [47], whereas others have described robust Ca^2+^ mobilization [48], emphasizing strong cell-type-specific FFA2 signaling. Similarly, 4-CMTB-mediated inhibition of GLP-1 release is consistent with studies reporting suppressed secretion in human and mouse FFA2-expressing islet cells at high concentrations [49]. Collectively, our findings suggest that FFA2 plays an inhibitory role in colonic GLP-1 releasing EECs, mediated through ligand bias and non-canonical signaling pathways.

The stimulatory role of FFA3 in colonic EECs was unexpected given that prior overexpression systems reported canonical Ga_i_-FFA3 coupling [20], which produced inhibitory outcomes. We found that AR420626-induced FFA3 activation promoted GLP-1 secretion via a Ca^2+-^dependent, PTX-sensitive pathway (Figures 5 and 6). Structural modeling has suggested that AR420626 targets a distinct allosteric pocket on FFA3 and directly interacts with Ga_i_ [50], a mechanism likely reflecting biased signaling at FFA3. We speculate that the βγ subunits released upon FFA3 activation stimulate PLCβ, leading to IP_3_-dependent Ca^2+^ mobilization, a mechanism previously described in other Ga_i_-coupled receptors such as a_2A_-adrenergic receptors [51]. Although this pathway is expected to produce only a modest rise in intracellular Ca^2+^ independent of Ga_q_ signaling [52], combined with other nutrient stimulation would be sufficient to drive GLP-1 exocytosis. This mechanism could explain how AR420626 stimulated GLP-1 release in primary mouse colonic cultures [16] and increased intracellular Ca^2+^ mobilization in FFA3-expressing cells, promoting glucose uptake [53]. Future work should resolve the structural basis of this non-canonical signaling, including how AR420626 stabilizes specific FFA3 conformations, and determine whether nutritional or metabolic changes alter these conformations in ways that modulate gut peptide expression [54], [55] and secretion [56], potentially impacting SCFA receptor signaling.

Given the structural similarities of ketone bodies to SCFAs (Figure 7A), our findings suggest that AcAc activates FFA2 to inhibit GLP-1 release in a cAMP-dependent manner, similar to the Ga_i_-biased ligand AZ1729 but in lower potency and consistent with effects reported in other FFA2-expressing systems [57]. BHB, in contrast, appears to act via FFA3, stimulating GLP-1 secretion through a yet-unidentified, non-cAMP, non-Ca^2+^ pathway (Figure 7B-I). This finding contrasts prior reports describing BHB as either an antagonist or inhibitory agonist of FFA3 [22], [21], [23], but supports our observations that both BHB and AR420626 can bias FFA3 and promote GLP-1 release in our system. The lack of Ca^2+^ mobilization by BHB likely reflects interaction of the orthosteric binding site rather than the positive allosteric site utilized by AR420626, suggesting that endogenous and synthetic ligands stabilize distinct FFA3 conformations toward a secretory outcome.

The ability of FFA2 and FFA3 to discriminate among SCFAs suggests a finely tuned ligand recognition mechanism that may extend to ketone bodies [50], potentially explaining their rank order of potency at each receptor. Structural modeling indicates that the binding cavity of FFA2 is smaller than that of FFA1, a receptor primarily responsive to medium- to long-chain fatty acids, whereas FFA3 is intermediate in size [58]. This geometry is consistent with FFA2 preferentially binding short SCFAs and FFA3 accommodating longer SCFAs. Within these pockets, conserved arginine residues (R180 and R255) are critical for receptor activation [59], and the predictive surrounding residues (Y90, I145 and E166 in FFA2, and F96, Y151 and L171 in FFA3) may impose steric constraints that confer chain length selectivity and ligand specificity [58]. These structural features likely reflect the preferential signaling of AcAc through FFA2 and BHB through FFA3: AcAc’s ketone position and carboxylate group are compatible with the tighter FFA2 pocket, while BHB’s additional hydroxyl and slightly longer chain are better accommodated by the larger FFA3 cavity. In contrast, smaller, more flexible SCFAs can access both binding pockets more promiscuously, as they are not constrained by rigid geometries. Ultimately, resolving the crystal structure of FFA3 will be essential to fully define the ligand-binding properties and downstream signaling mechanisms.

## 5. Conclusions

Our findings reveal that FFA2 and FFA3 in colonic GLP-1 releasing EECs respond to SCFAs and ketone bodies through distinct, non-canonical pathways, enabling ligand-dependent regulation of gut hormone secretion. This flexibility allows GLP-1 release to adapt to nutrient availability, increasing after carbohydrate intake while remaining responsive under low-carbohydrate conditions through ketone body activation. Together, these results highlight FFA2 and FFA3 as key integrators of metabolic signals, fine-tuning EEC function to dietary and metabolic cues. Future work should define the structural basis of ligand recognition and explore how these mechanisms are altered in metabolic disease to guide therapeutic strategies.

## CRediT authorship contribution statement

**Karly E. Masse:** Writing – review & editing, Writing – original draft, Visualization, Validation, Methodology, Investigation, Formal analysis, Conceptualization. **Brandon Lee:** Methodology, Investigation. **HengLiang Wu:** Methodology, Resources. **Jackie Han:** Methodology. **Pierre Larraufie:** Writing – review & editing, Methodology, Resources. **Frank Reimann:** Writing – review & editing, Resources. **Fiona M. Gribble:** Writing – review & editing, Resources. **Van B. Lu:** Writing – review & editing, Validation, Supervision, Resources, Project administration, Funding acquisition, Data curation, Conceptualization.

## Acknowledgments

We thank Daniel J. Drucker (Toronto) for permission to use the GLUTag cell line.

## Glossary

AcAc: Acetoacetate
BBS: Bombesin
BHB: β-hydroxybutyrate
C2: Acetate
C3: Propionate
C4: Butyrate
EEC: Enteroendocrine cell
FFA1: Free fatty acid receptor 1
FFA2: Free fatty acid receptor 2
FFA3: Free fatty acid receptor 3
GLP-1: Glucagon-like peptide-1
GPCR: G-protein coupled receptor
OZITX: Ga_o_, Ga_z_, Ga_i_ inhibiting toxin
PTX: Pertussis toxin
SCFA: Short-chain fatty acid
SST: Somatostatin

