## Supplemental files for "Non-canonical signaling mechanisms of short-chain fatty acid receptors in glucagon-like peptide-1 (GLP-1) releasing enteroendocrine cells"

### Non-canonical signaling mechanisms of SCFA receptors in GLP-1 releasing EECs

KE Masse et al.

Submitted to Molecular Metabolism

Supplementary Figures

#### Suppl Figure 1:

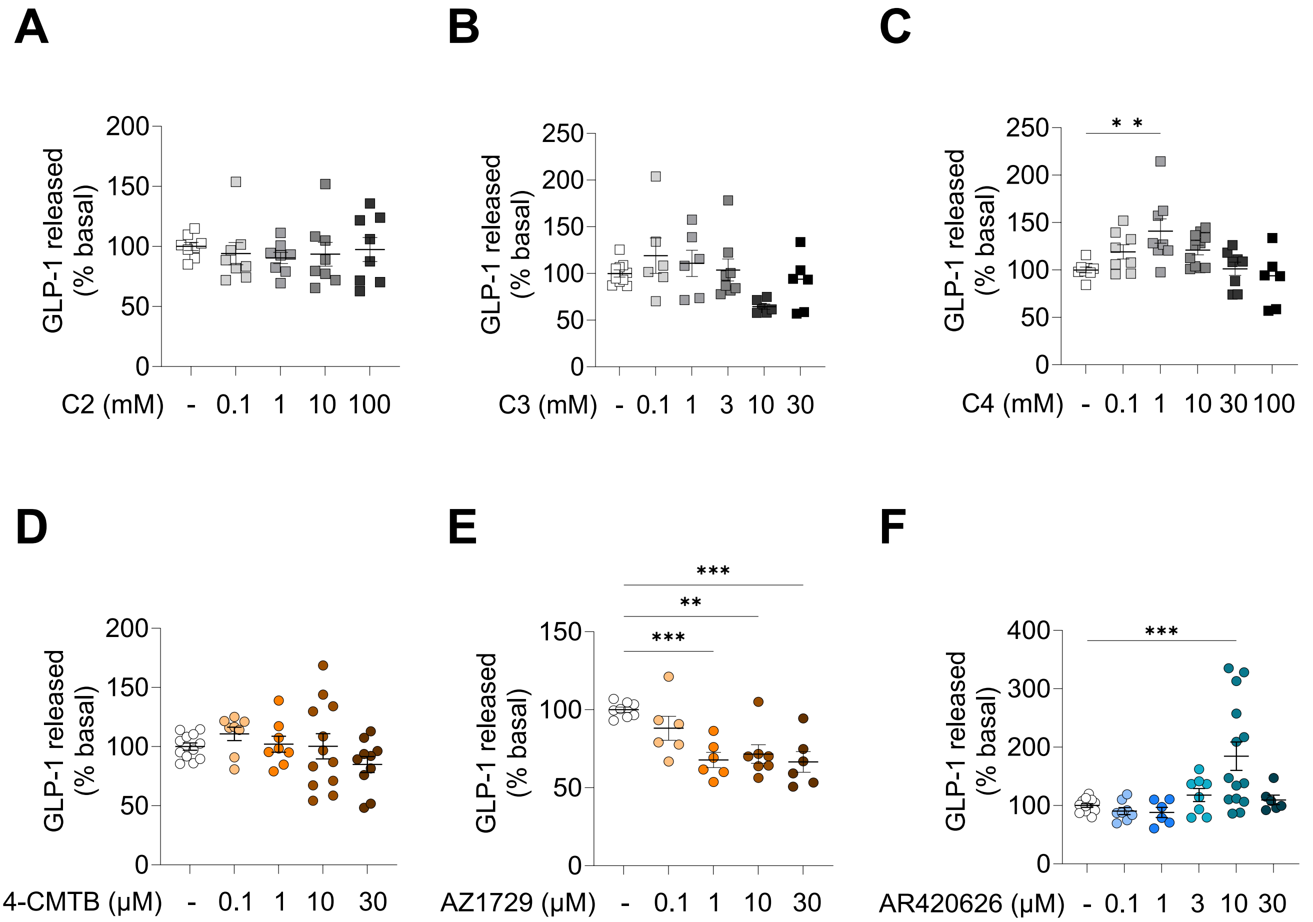

### Suppl Figure 2:

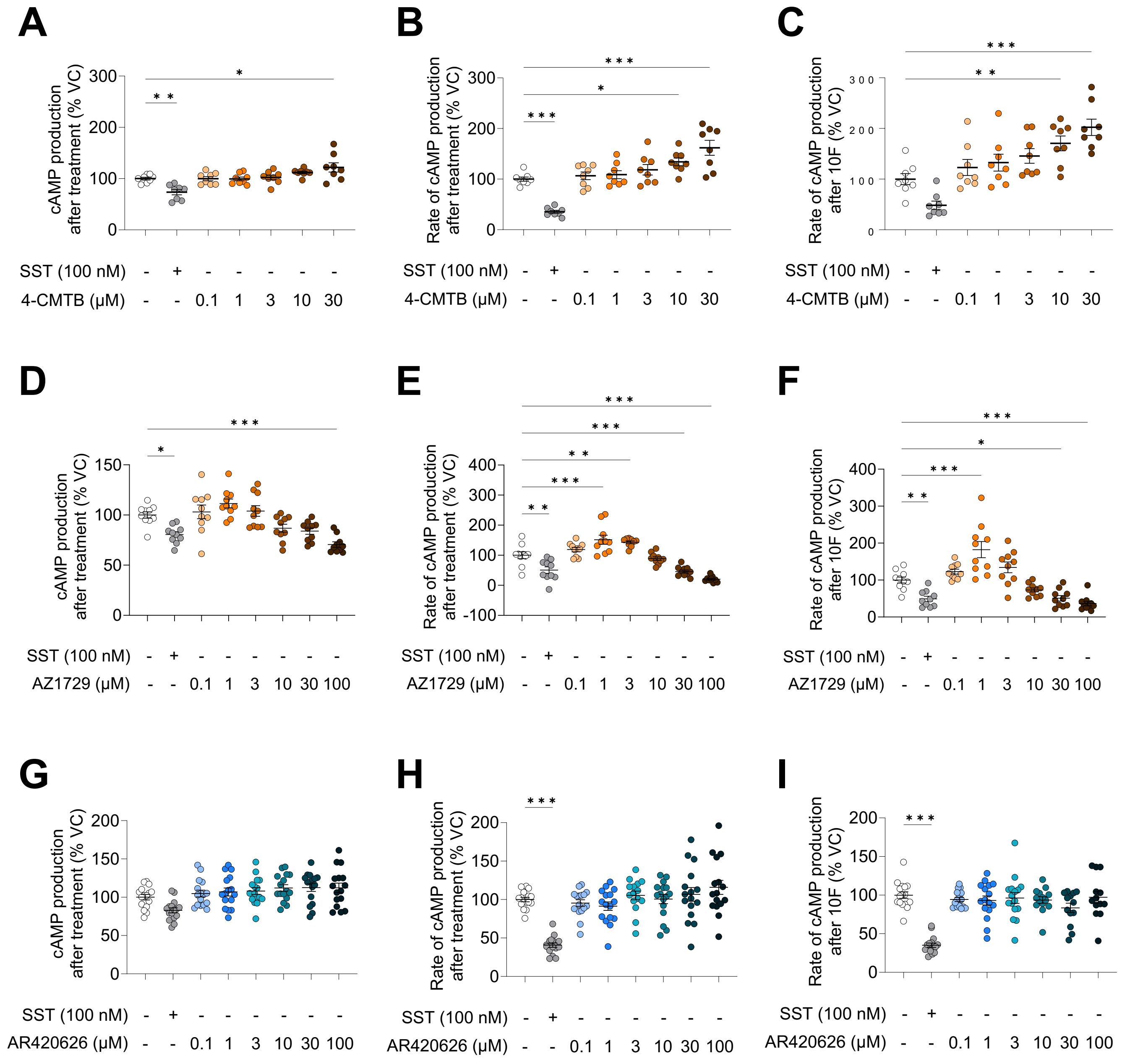

### Suppl Figure 3:

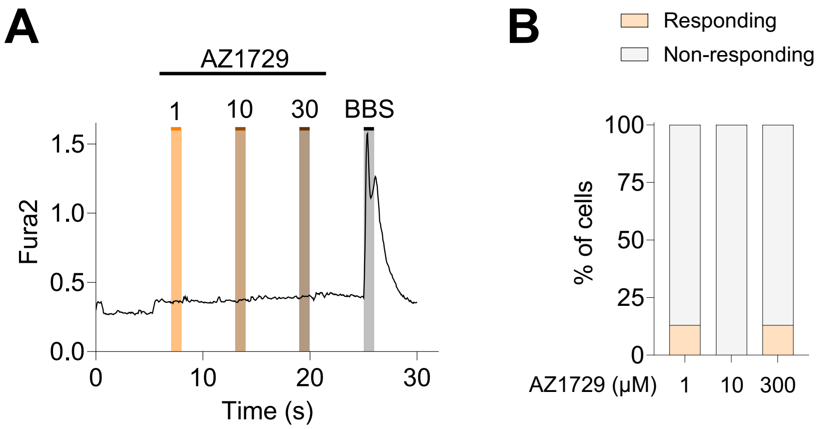

#### Suppl Figure 4:

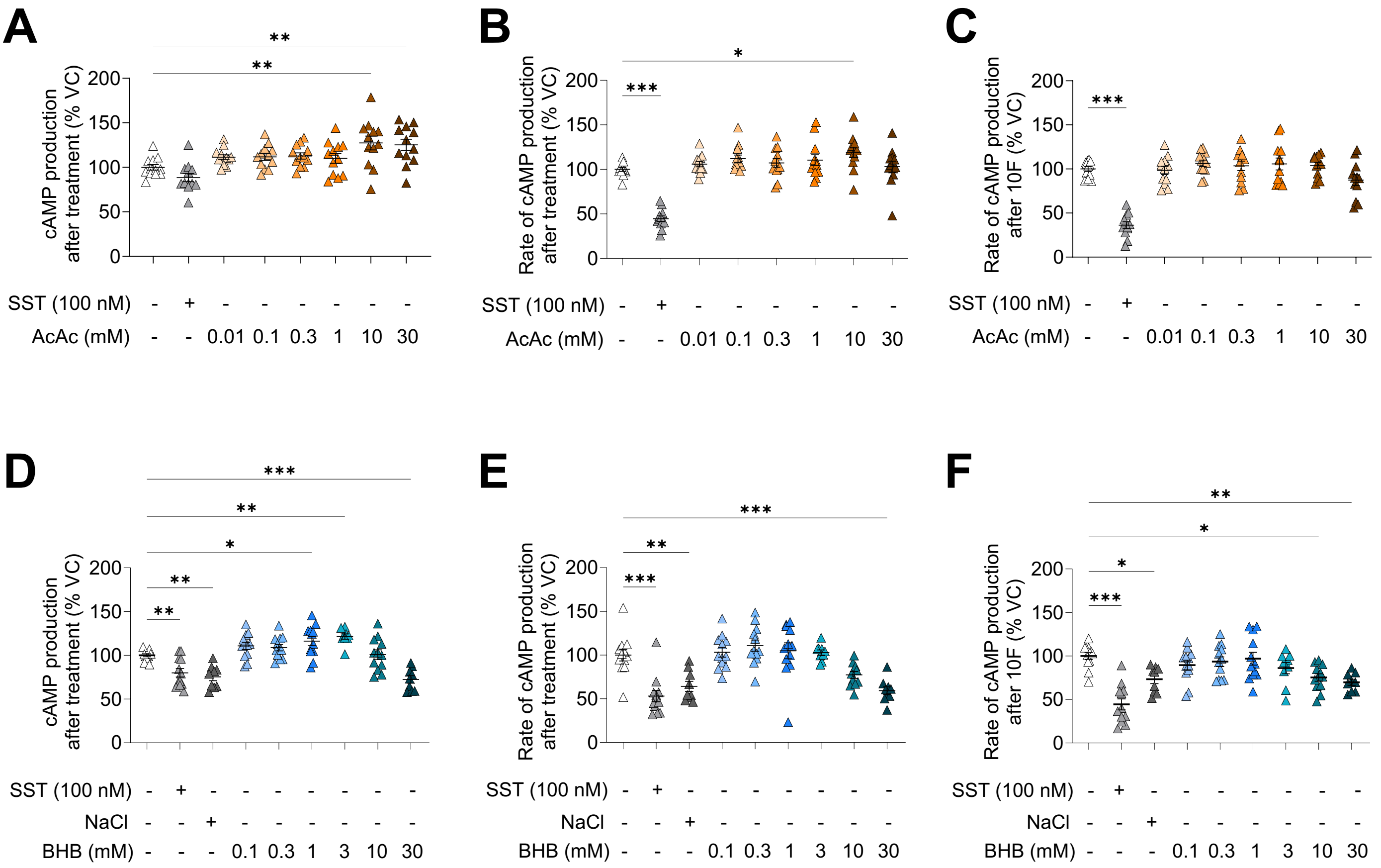

Supplementary Figure Legends:

Supplementary Figure 1: **Effects of SCFAs and select FFA2/3 ligands on basal GLP-1 secretion in GLUTag cells.**

GLP-1 levels released from GLUTag cells following 2h treatment with a standard saline solution and the addition of individual SCFAs including acetate (A), propionate (B) or butyrate (C). Similar secretion experiments testing 2h treatment with a standard saline solution and a selective ligand of free fatty acid receptor 2 (FFA2), 4-CMTB (D) or AZ1729 (E), or free fatty acid receptor 3 (FFA3), AR420626 (F). All secretion data was normalized to total protein from cell lysates and expressed relative to the standard saline solution alone condition. Individual symbols represent GLP-1 released per well. Orange symbols represent responses targeting FFA2, blue symbols represent responses targeting FFA3, grey symbols represent potential activation of multiple SCFA-sensitive receptors. N = 6-18 from at least 3 independent experiments, mean ± SEM indicated. One-way ANOVA with Dunnett's multiple comparisons statistical test performed; **P<0.05, **P<0.01, ***P<0.001*.

Supplementary Figure 2: **Changes in intracellular cAMP production following treatment with select FFA2/3 ligands in GLUTag cells stably expressing GloSensor-22F™.**

(A, D, G) Calculated intracellular cAMP production after treatment. Determined as the maximal response immediately following addition of select ligands, normalized to the maximal response of vehicle control. (B, E, H) Calculated rate of cAMP production after treatment. Determined as the maximum slope of intracellular cAMP production immediately following addition of select ligands, normalized as a fraction of the maximal slope response of vehicle control. (C, F, I) Calculated rate of cAMP production after forskolin treatment. Determined as the maximum slope of intracellular cAMP production during the application of forskolin for each select ligand, normalized as a fraction of the maximal slope response during the application of forskolin of vehicle control. Application of the select FFA2 ligand 4-CMTB (A-C) or AZ1729 (D-F), and FFA3 ligand AR420626 (G-I) were tested. Individual symbols represent a single well. N = 8-16 from at least 3 independent experiments, mean ± SEM indicated. One-way ANOVA with Dunnett's multiple comparisons statistical test performed; **P<0.05, **P<0.01, ***P<0.001*.

Supplementary Figure 3: **No intracellular $Ca^{2+}$ response to AZ1729 in GLUTag cells.**

Representative recording of intracellular $Ca^{2+}$ levels during treatment with AZ1729 (1 – 30 µM) and 100 nM bombesin (BBS) (A). Intracellular $Ca^{2+}$ measured using ratiometric Fura2 fluorescence at 340 and 380 nm excitation. Drugs were applied as indicated above trace. Calculated % of GLUTag cells responding to AZ1729 treatment with an increase in Fura2 fluorescence above a pre-set threshold (>0.05) (E). Orange bar indicates % of cells that responded to AZ1729 treatment, grey bar indicates % of cells that did not respond to AZ1729 treatment. N = 8 from at least 3 independent experiments.

Supplementary Figure 4: **Changes in intracellular cAMP production following treatment with ketone bodies in GLUTag cells staby expressing GloSensor-22F™.**

(A, D) Calculated intracellular cAMP production after treatment. Determined as the maximal response immediately following addition of select ligands, normalized to the maximal response of vehicle control. (B, E) Calculated rate of cAMP production after treatment. Determined as the maximum slope of intracellular cAMP production immediately following addition of select ligands, normalized as a fraction of the maximal slope response of vehicle control. (C, F) Calculated rate of cAMP production after forskolin treatment. Determined as the maximum slope of intracellular cAMP production during the application of forskolin for each select ligand, normalized as a fraction of the maximal slope response during the application of forskolin of vehicle control. Application of acetoacetate, AcAc (A-C) or β–hydroxybutyrate, BHB (D-F) were tested. Individual symbols represent a single well. N = 9-12 from at least 3 independent experiments, mean ± SEM indicated. One-way ANOVA with Dunnett's multiple comparisons statistical test performed; **P<0.05, **P<0.01, ***P<0.001*.
